# Calibration improves estimation of linkage disequilibrium on low sample sizes

**DOI:** 10.64898/2026.03.05.709321

**Authors:** Ulises Bercovich, Carsten Wiuf, Anders Albrechtsen

## Abstract

Linkage disequilibrium is a central statistic in population genetic studies, commonly measured by the squared correlation between pairs of genetic variants. An important drawback of this measure is its upward bias caused by a finite sample size. To handle this, different methods exist that correct for sample-size bias. However, because the correlation consists of a ratio, there is no unbiased method to compute it. In this work, we present a procedure to calibrate those methods using a non-parametric approach with simulated data. This is done with forward modeling to generate genotype matrices with known parameters, followed by an inverse mapping to recover estimates of the underlying parameters. Then, a mean-centering calibration is applied to the recovered estimate of the true parameter. This approach is applied to real and simulated data, showing consistent improvement in accuracy compared to other sample-size-aware methods. Furthermore, to study the effects on downstream analyses, we analyze the classification performance on LD pruning, where we also observe an improvement, particularly in extreme cases with low sample sizes of 5 or 10 individuals.

## Introduction

Linkage disequilibrium (LD) is a measure of non-random association between alleles at different loci. The structure of LD in a population is shaped by different evolutionary forces: it is primarily generated by genetic drift and broken down by recombination, but is also influenced by mutation, population size changes, and admixture, among other forces [1, 2]. Hence, by understanding how these forces shape LD, valuable inferences can be made by studying the LD structure of a population, such as reconstructing its demographic history or identifying selection [3, 4]. Moreover, since LD violates the assumption of independence between genetic markers, accounting for its structure is necessary for a wide range of methods that assume independence between variants.

LD is commonly calculated as the squared sample correlation coefficient among genetic markers [1]. This classical estimate is an effective measure for genetic association in large samples. It is based on the sample covariance, an unbiased estimator of the covariance. However, because the correlation consists of the ratio of the estimated covariance to the product of the variances, it does not inherit the unbiasedness of its components [5]. This bias is even stronger when considering the square of the sample correlation. In particular, for low sample sizes, there is an upward bias that can affect different LD-dependent methods, such as LD curves and LD pruning [6]. Furthermore, downstream analyses based on pruned data, such as the fixation index or principal component analysis, are also affected by this bias solely generated by the sample size.

This problem is especially relevant in population genetic studies where small sample sizes are common. In conservation and studies of rare or remote species, sample collection can be a major challenge. Similarly, in ancient DNA studies samples may be sparse without the possibility of extension. Even in the analyses of contemporary human data, targeting a specific subpopulation may result in insufficient sample sizes. In these contexts, poor estimates of LD can lead to incorrect inference, and increasing the sample size is often impossible, so statistical correction is the only option.

Numerous studies have proposed methods to address the sample size bias of correlation coefficients between normally distributed variables [7, 8]. In these cases, an approximation of the density of the Pearson correlation (*r*) can be calculated analytically based on properties of normal densities [9]. This proved to be helpful in constructing corrections for the bias that are most notorious when dealing with low sample sizes. However, no single correction performs best across all settings, and these methods rely on assumptions that do not translate to genotype data. More specifically, a major issue is that genomic data are discrete. For example, biallelic data is usually described using a matrix of 0s and 1s for phased data, or 0s, 1s, and 2s for unphased data. Hence, the knowledge of an approximation of the density in the normal case cannot be transferred to the binomial or multinomial distributions of genomic data.

Deriving the true probability density function for the correlation coefficient of binomial data is analytically intractable [10]. This is due to the discrete nature of the genotypes, which creates a prohibitive combinatorial problem when enumerating all possible sample configurations. The discrete and bounded nature of genomic data combined with low sample size, means that the problem does not allow the use of the Central Limit Theorem to have an approximation of the density. Therefore, asymptotic theory cannot be applied and specialized methods are required for genomic data [5, 11].

In this work, we introduce a model-free two-step calibration procedure designed to correct the bias of LD estimates at small sample sizes. While the study of sampling distributions of statistics in population genetics is not new [12], their use has traditionally been descriptive rather than corrective [13]. Here, we exploit these distributions to directly calibrate the estimation of LD. The first step relies on simulation: we generate multiple genotype matrices under known parameters and record their LD estimates. This simulation grid is then used in an inverse regression framework, allowing us to map any observed estimate back to the most likely underlying parameter value. The second step refines this correction by adjusting the mean, following the structure of known sample-size corrections. This additional calibration is especially valuable for settings where the mean is critical, such as LD curves, where reducing bias is worth a moderate increase in variance. This two-step procedure can be applied not only to the uncorrected squared correlation coefficient but also to existing sample-size-aware estimators.

In this project, we evaluate the calibration procedure on the classical estimator of the squared correlation. This is compared to three correction methods, including a novel approach that emphasizes stability and potential applicability to private data, following the ideas of [5]. To study the effects of calibration, we use real data from the 1000 Genomes Project [14] and simulated data based on an African population [15] using stdpopsim [16]. We then apply the calibration procedure to the analyzed method, where we consistently observe improvement in the mean square error. Furthermore, this improvement is passed on to LD pruning, where we measure the classification performance of the different approaches.

## Methods

Let *G* be a genotype matrix of size *n*×*m*, where *n* is the number of individuals and *m* the number of variants. In this work we focus on the small-sample setting (*n* < 50), although the method can be extended to any *n*. We assume that *G* represents unphased data, though it generalizes naturally to phased genotypes. For an individual *i* at locus *s*, the marginal distribution of *G*_*is*_ is Bin(2, *p*_*s*_), where *p*_*s*_ denotes the allele frequency at that site. To quantify LD between two loci *s* and *t*, let *p*_*st*_ denote the probability that both loci carry the reference allele on a haplotype. Then the covariance between these loci satisfies

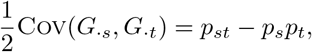

as shown in [17]. Denote *D*_*st*_:= Cov(*G*_·*s*_, *G*_·*t*_)/2. The population squared correlation is defined as

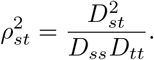

In practice, 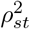 is estimated by the squared sample correlation,

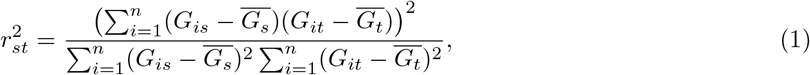

where 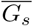 is the sample mean at locus *s*. This estimator is consistent as *n* → ∞, but it is biased upward for small *n* [6]. For example, if loci *s* and *t* are truly independent, then 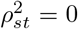, whereas the non-negativity of 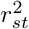 implies 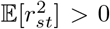. This bias is most pronounced when sample sizes are very small and when the true correlation is close to zero.

In the low-sample-size setting, we use forward modeling to generate genotype matrices and study how sample size affects the squared correlation. Specifically, fix *n* and consider a valid tuple 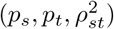. Here, a tuple is considered valid if the covariance inherited by these parameters, which is

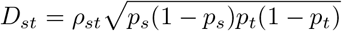

(up to a sign difference) is within the range

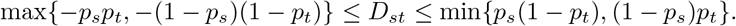

This ensures that the corresponding haplotype frequencies are between 0 and 1 [18]. Based on these parameters, we generate random samples of the form 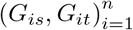 with *G*_*is*_ ∼ Bin(2, *p*_*s*_), *G*_*it*_ ∼ Bin(2, *p*_*t*_), and correlation 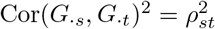. For each simulated sample, we compute 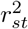, which is then used to compute an average over the replicates. This mapping between the observed average of 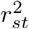 and the true 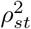 then allows us to perform inverse regression, yielding a calibrated estimator of the population squared correlation.

To formalize the approach, consider a valid tuple 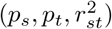, and define

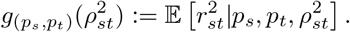

Intuitively, the function *g* describes the expected distortion introduced by finite sample size. It maps the true population correlation 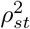 to the average value of the statistic based on the sample 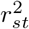. For fixed (*p*_*s*_, *p*_*t*_), this mapping is increasing in 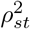: higher true correlation produces, on average, higher observed *r*^2^. Hence, its inverse exists and can be used to de-bias the observed statistic. Therefore, we calculate the inverse function to recover the true 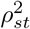. That is,

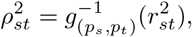

where 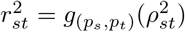. Then, we define our calibrated estimator as

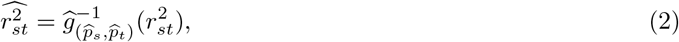

where 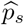 and 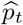 are allele-frequency estimators.

In Figure 1, we illustrate the relationship between the true squared correlation 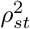 and the expected sample statistic, 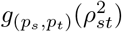. For three different sample sizes (*n* ∈ *{*5, 10, 25*}*) and fixed population allele frequencies (*p*_*s*_, *p*_*t*_) = (0.5, 0.5), we simulated 5,000 genotype matrices for each value of 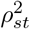 on a grid from 0 to 1 with a step size of 0.05. This was done by first calculating the haplotype frequencies that correspond to the tuple 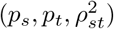 and then sampling genotypes for the from these assuming random mating. For each replicate, we computed the observed 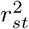 and then averaged across replicates. The resulting estimated 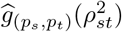 is shown in blue. The red diagonal corresponds to the identity line, representing an unbiased estimator. The vertical distance between the blue curve and the diagonal therefore quantifies the bias. Notably, the upward bias is strongest for small *n* and for true correlations close to zero.

**Figure 1:**
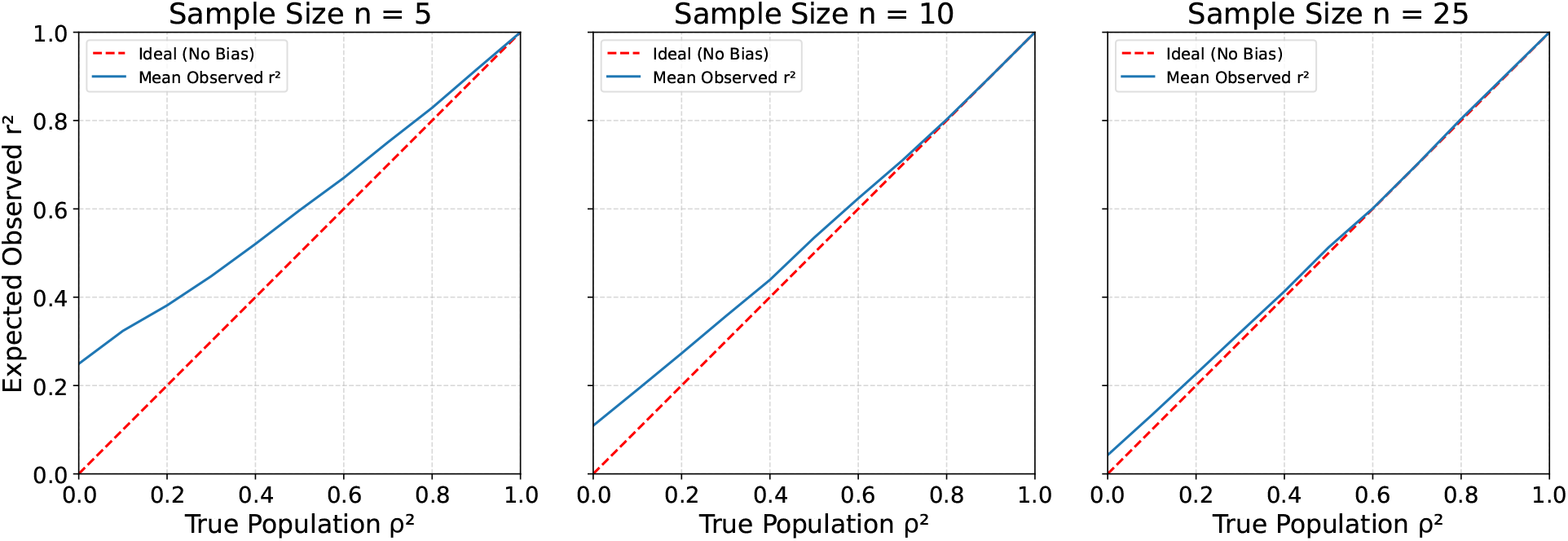
Bias curves for the sample *r*^2^ estimator across different sample sizes with fixed allele frequencies of (*p*_*s*_, *p*_*t*_) = (0.5, 0.5). Each panel plots the true population *ρ*^2^ (x-axis) against the mean observed 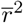 (y-axis) from 5,000 simulations. The red diagonal represents the line of an ideal, unbiased estimator. Moving across the columns (n = 5, 10, 25) shows that the upward bias is progressively reduced with larger sample sizes.

In practice, calibration is performed by precomputing these bias curves for all possible sample allele-frequency pairs 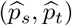, of which there are (2*n*)^2^ in the unphased diploid setting. Given an observed geno-type matrix, the empirical allele frequencies 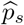 and 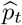 are obtained via 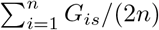 and 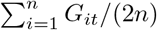, respectively. The observed 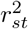 is then de-biased by mapping it through the inverse of the precomputed curve for 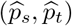. For example, with *n* = 5 individuals and 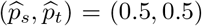, Figure 1 shows that an observed value of 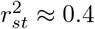 corresponds to a calibrated estimate 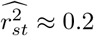. The algorithm describing the procedure to estimate the map 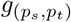 and its use for calibration can be found in Algorithm S2.

Although this approach may appear computationally intensive and unsuitable for large genotype datasets, the bias curves can be generated once in advance, requiring only seconds to minutes depending on *n*. Additionally, in the low-sample-size regime, the space of possible values of 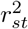 is discrete and relatively small, enabling efficient precomputation. Thus, the calibration step itself reduces to a simple table lookup, adding negligible runtime overhead when applied to real data.

This calibration framework can be directly extended to estimators that already incorporate sample-size corrections for LD. For example, instead of starting from 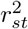, one may consider alternatives proposed in the literature [5, 6, 11]. Applying the same procedure yields new calibration curves that map the sample-size–corrected statistics to their population counterparts. An important advantage of this approach is that it corrects for the well-known issue that some sample-size–aware estimators may fall outside the admissible range [0, 1]. A study of this can be found in the Supplementary Section S2.

While this non-parametric calibration effectively reduces the upward bias induced by small sample sizes, constraining the estimators to the [0, 1] interval inevitably leaves a residual upward bias near zero. To address this, we introduce a second calibration step that allows estimators to take negative values. This step again relies on forward modeling but is designed to mean-center the distribution of the calibrated estimator under independence. Such a correction is particularly relevant in applications where the shape of LD decay curves matters, because bias near the lower tail can distort interpretation.

The idea is motivated by the algebraic form of existing corrections [11, 19, 20], which can be expressed as

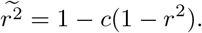

By focusing on the case of independent pairs 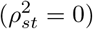, we estimate values 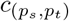 such that

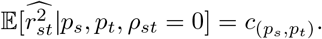

The second calibration step is then defined as

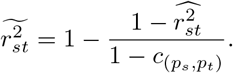

By construction, when 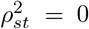 we obtain 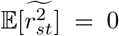, thus ensuring unbiasedness under independence. Allowing negative values in this controlled way provides an estimator with correct behavior at the lower tail of LD curves, supported by both theory and simulation.

## Methods to correct linkage disequilibrium used for comparison

To evaluate the benefits of the proposed two-step calibration on the squared correlation, we applied it to several existing estimators of LD, along with the approach introduced in this work. The first is the correction by Bulik-Sullivan et al. [11],

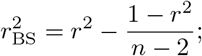

which adjusts the naive *r*^2^ by reducing the upward bias induced by small sample sizes. The second is the estimator by Ragsdale and Gravel [5], denoted 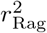, which relies on unbiased estimators of the numerator and denominator of *r*^2^ to provide a more stable correction across sample sizes. A third sample-size-aware method, denoted 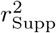, is described in Supplementary Material S1.

### Data

The performance of these estimators was assessed using two datasets: one based on real human data and one generated by coalescent simulations. In both cases, we focused on high-quality unphased SNP data. The real dataset was obtained from the 1000 Genomes Project [14], and consists of 378 Utah residents with Northern and Western European ancestry (CEU). Only variants from chromosome 22 were considered, further restricted to SNPs from the HapMap3 panel [21]. After applying a minor allele frequency (MAF) filter of 5%, the dataset comprised 14,465 SNPs. This dataset is available via the `magenpy` software [22].

The simulated dataset was generated with the `stdpopsim` package [16, 23], which standardizes demographic models for population genetic simulations using `msprime` [24] and `SLiM` [25]. We followed the African demographic model (ID: Africa 1T12) described in [15]. A total of 400 individuals were simulated for chromosome 22, restricted to the 20–30 Mb region. After applying the same MAF filter of 5%,, the resulting dataset contained 13,543 SNPs.

### Metrics

To measure the accuracy of each method, we use the root mean square error (RMSE), which is given by

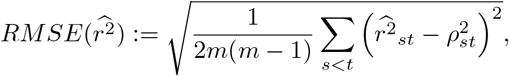

where *m* is the number of SNPs in the data. This measures the average deviation of the estimated squared correlation from its true population value.

To measure the effects of calibration on LD pruning, we quantify the performance of the methods with the F1 score. This is a standard metric for classification that balances true positives (TP), false positives (FP), true negatives (TN) and false negatives (FN). By counting these groups, we calculate F1 score with the harmonic mean of the metrics Precision and Recall. In this context, a true positive is a pair of SNPs in high LD where pruning correctly removes one variant. Precision, defined as TP/(TP+FP), is related to *over-pruning*; since a low Precision is equivalent to having many pairs of variants that are incorrectly flagged as dependent and removed (false positives). Recall, defined as TP/(TP+FN), quantifies *under-pruning*, as a low recall indicates that a large number of kept pairs that are dependent (false negatives).

For both RMSE and F1, the ground truth is approximated by the squared correlation estimated from the full dataset. To avoid bias toward distant SNP pairs, which are far more numerous and typically independent, we stratify pairs by physical distance into bins. Each bin corresponds to an interval of base-pair distances, and within each bin we select an equal number of pairs. This ensures that the error metrics reflect performance across different genomic scales, rather than being dominated by the abundance of long-range independent pairs.

## Results

To evaluate the performance of our proposed estimators, we conducted a series of bootstrap experiments using two distinct datasets. The first is a real-world dataset (CEU) consisting of *n*=378 individuals of European ancestry from the 1000 Genomes Project [14]. The second is a simulated dataset (AFR) generated with `stdpopsim` [16] using a standard African demographic model with *n*=400 individuals, providing a distinct LD structure. For our benchmark, we compared the standard sample *r*^2^ (“Samp”) and three previously mentioned corrected estimators (“BS”, “Rag”, “Supp”) against our two novel proposals: a primary calibrated estimator (“Cal”) and a two-step calibration (“mCal”), where we do a further mean-correction on the “Cal” estimator. For the calibration procedures, 5000 genotypes matrix were generated for each tuple (*p*_*s*_, *p*_*t*_, *ρ*^2^). The *p*_*s*_ and *p*_*t*_ values were computed in a grid from 0.02 to 0.5 with a step of 0.02. For *ρ*^2^, the grid was from 0 to 1 in 20 steps, considering that for most of the pairs (*p*_*s*_, *p*_*t*_), there is an upper bound for the maximum achievable *ρ*^2^ which is lower than 1.

To assess the accuracy and stability of these estimators, we designed a unified bootstrap procedure that was applied to generate results for both the RMSE and the F1 score. This procedure was repeated 100 times for each sample size (*n* = 5, 10, 25). In each of the 100 replicates, we first drew a random subsample of *n* individuals from the full dataset. Then, we filtered the sub-sampled genotype matrix to retain only polymorphic variants. From this set of valid SNPs, a new, random set of SNP pairs was selected using a bin-stratified sampling approach, where up to 1,000 pairs were drawn from each genomic distance bin (up to 1 Mbp). For this fixed set of SNP pairs, we then established the “ground truth” *ρ*^2^ by applying the standard sample *r*^2^ estimator to the genotypes from the full, original dataset. The *r*^2^ was also calculated for the same set of SNP pairs using the genotypes from the small *n*-individual subsample with each estimator.

The results for this experiment are presented in Figure 2. Here, we can see that the calibrated estimator “Cal” consistently has a higher accuracy than the other methods. Moreover, we can observe the effect of the two-step calibration that focuses on adjusting solely the bias, where we have an increment in the RMSE compared to the one-step calibration. In contrast, the bias is reduced, as it can be seen in the decomposition of the RMSE found in Supplementary Figures S4 and S5. A further application of having a reduction of bias while increasing the RMSE can be found in the Supplementary Section S3, where we show an improvement when estimating LD scores.

**Figure 2:**
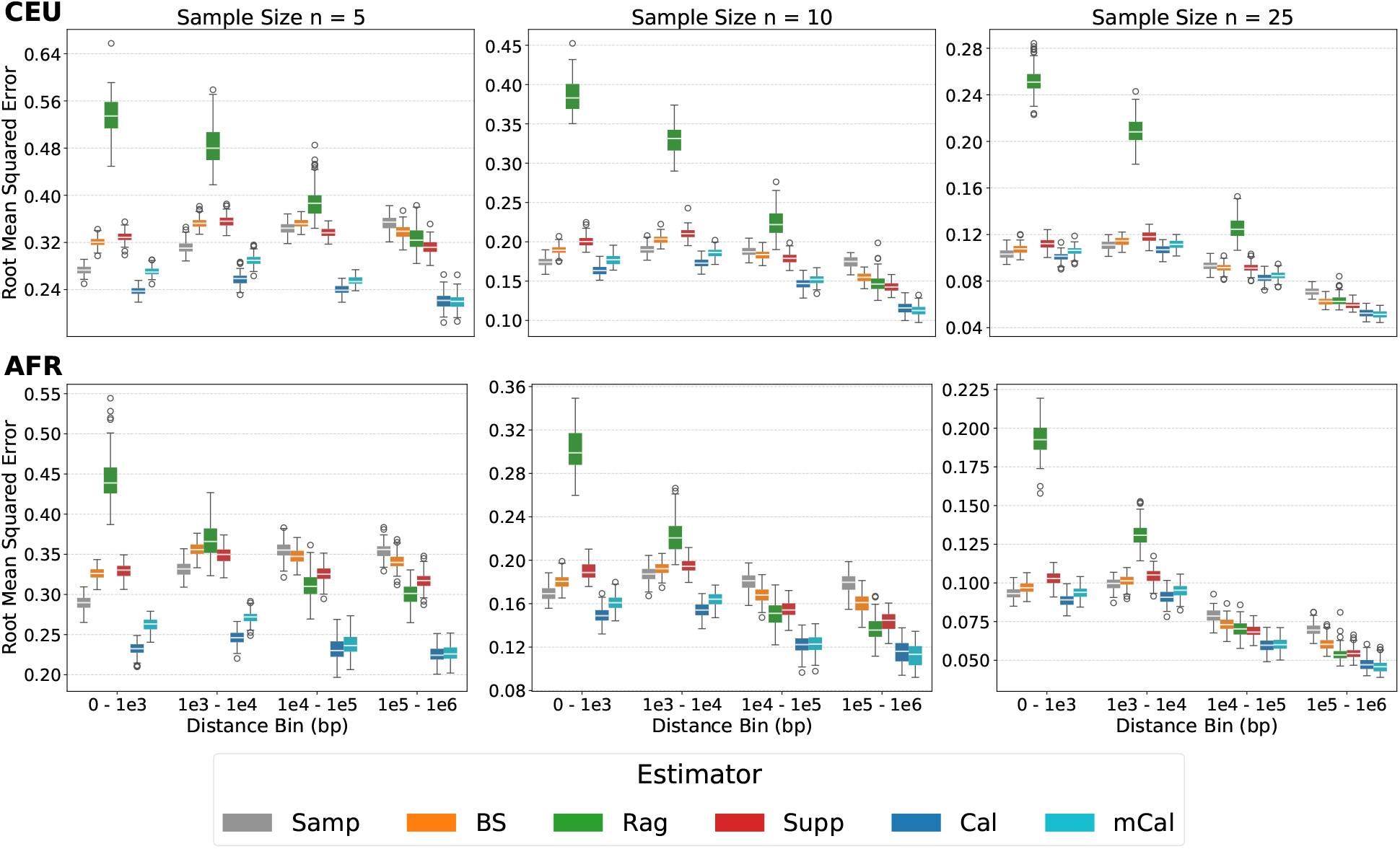
Distribution of RMSE for *r*^2^ estimators across genomic distance and sample size. Each row is computed by using different data (CEU and AFR), while each of the three columns corresponds to a different subsample size (*n*). The x-axis in each plot represents genomic distance, divided into bins. Within each bin, the boxplots show the distribution of RMSE values obtained from 100 independent bootstrap replicates, where each color represents a different *r*^2^ estimator.

To study the effects of pruning, we use the same as before but evaluate whether each pair of SNPs exceeds a certain LD threshold. We compare this to the results from using the population *r*^2^ to calculate F1 score. To perform a balanced computation of the metric, as before we stratified the pairs within bins and computed up to 1000 pairs within each one. This is to avoid the over representation of independent variants. Furthermore, we only computed pairs up to 1M basepairs (bp). In Figure 3, we observe how the calibrated methods outperform the uncalibrated methods, as a higher F1 score implies a better classification performance. This can be clearly seen in the cases with the lowest sample sizes of *n* = 5 and *n* = 10. Therefore, we observe an immediate effect of having increasing accuracy of estimating *r*^2^: LD pruning gets better.

**Figure 3:**
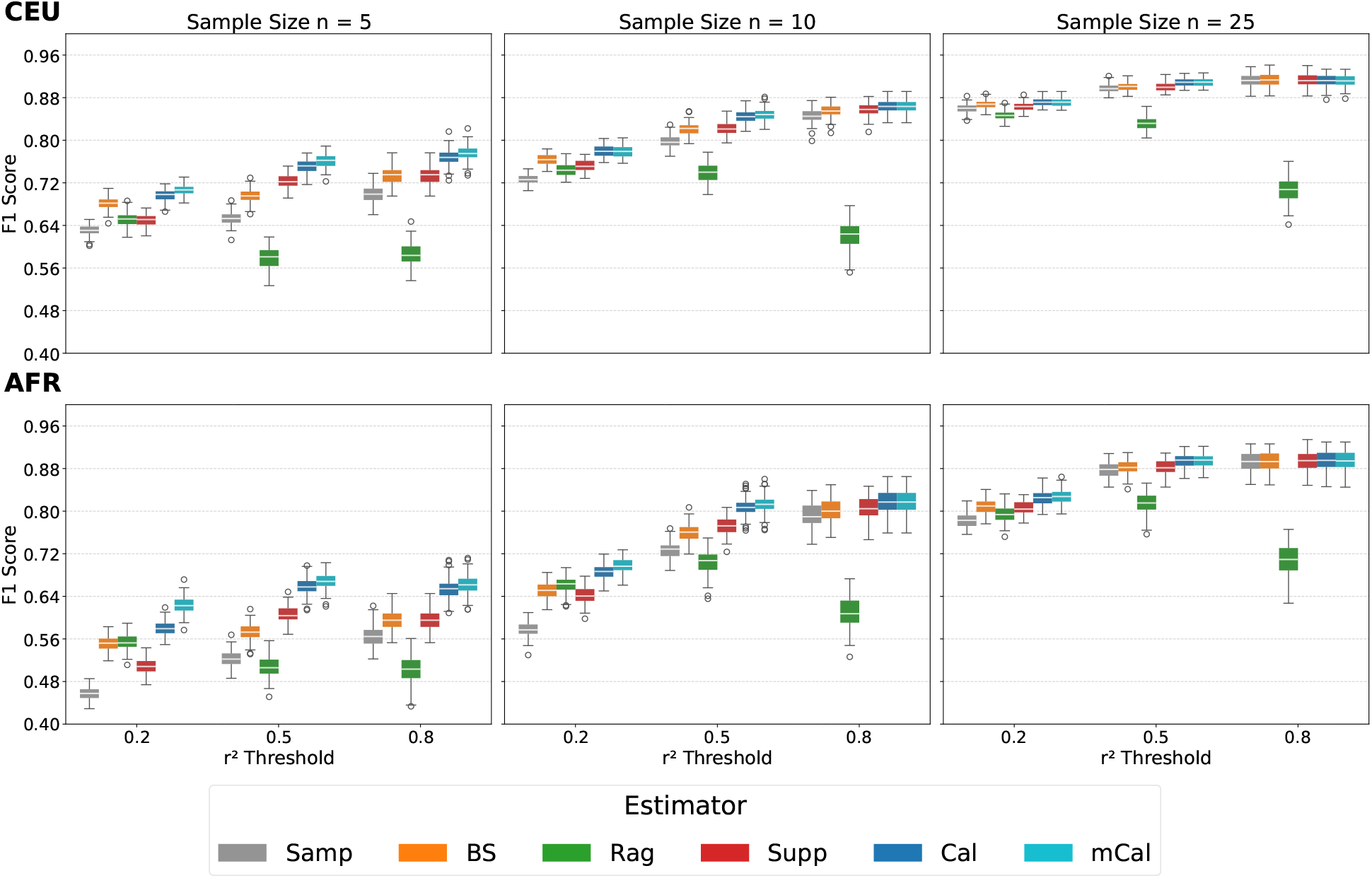
F1 Score for LD pruning performance, compared across populations and sample sizes. The top and bottom rows show performance on data from the CEU and AFR populations, respectively. Each column displays results for a different subsample size (*n*). The x-axis represents the *r*^2^ threshold for pruning, while the y-axis shows the F1 score, where a higher value indicates a better balance between precision (avoiding over-pruning) and recall (avoiding under-pruning). The boxplots are based on the resulting F1 score for each method over 100 independent bootstrap replicates.

We further investigated pruning performance by applying a simple, window-based pruning procedure to both datasets. For each SNP *s*, the estimated *r*^2^ against all other variants within 1 Mbp was computed. Any SNP *t* with *r*^2^ exceeding a predefined threshold (*r*^2^ = 0.2) was pruned. The process then moved to the next non-pruned variant and repeated until all variants were evaluated. To assess pruning quality, we measured the true *r*^2^ between adjacent retained SNPs and calculated the percentage below the threshold, along with the total number of variants kept. This approach, analogous to the F1 score, balances retaining more variants against avoiding under-pruning.

Figure 4 illustrates pruning outcomes for *n* = 10 (*n* = 5 and *n* = 25 are shown in Supplementary Figures S6 and S7). In both the CEU and AFR datasets, most adjacent SNP pairs have low LD, close to zero. However, the number of retained variants and pruning efficiency vary by method. The Ragsdale estimator (“Rag”) consistently retains the largest number of variants, but at the cost of higher under-pruning. In contrast, the standard sample *r*^2^ (“Samp”) yields the lowest misclassification but keeps less than half of the variants compared to other methods. BS and Supp estimators fall in between. The calibrated methods (“Cal” and “mCal”) achieve a favorable balance, maintaining low misclassification while retaining more variants. This illustrates why the calibration procedures achieve higher F1 scores: they effectively balance over- and under-pruning to optimize performance.

**Figure 4:**
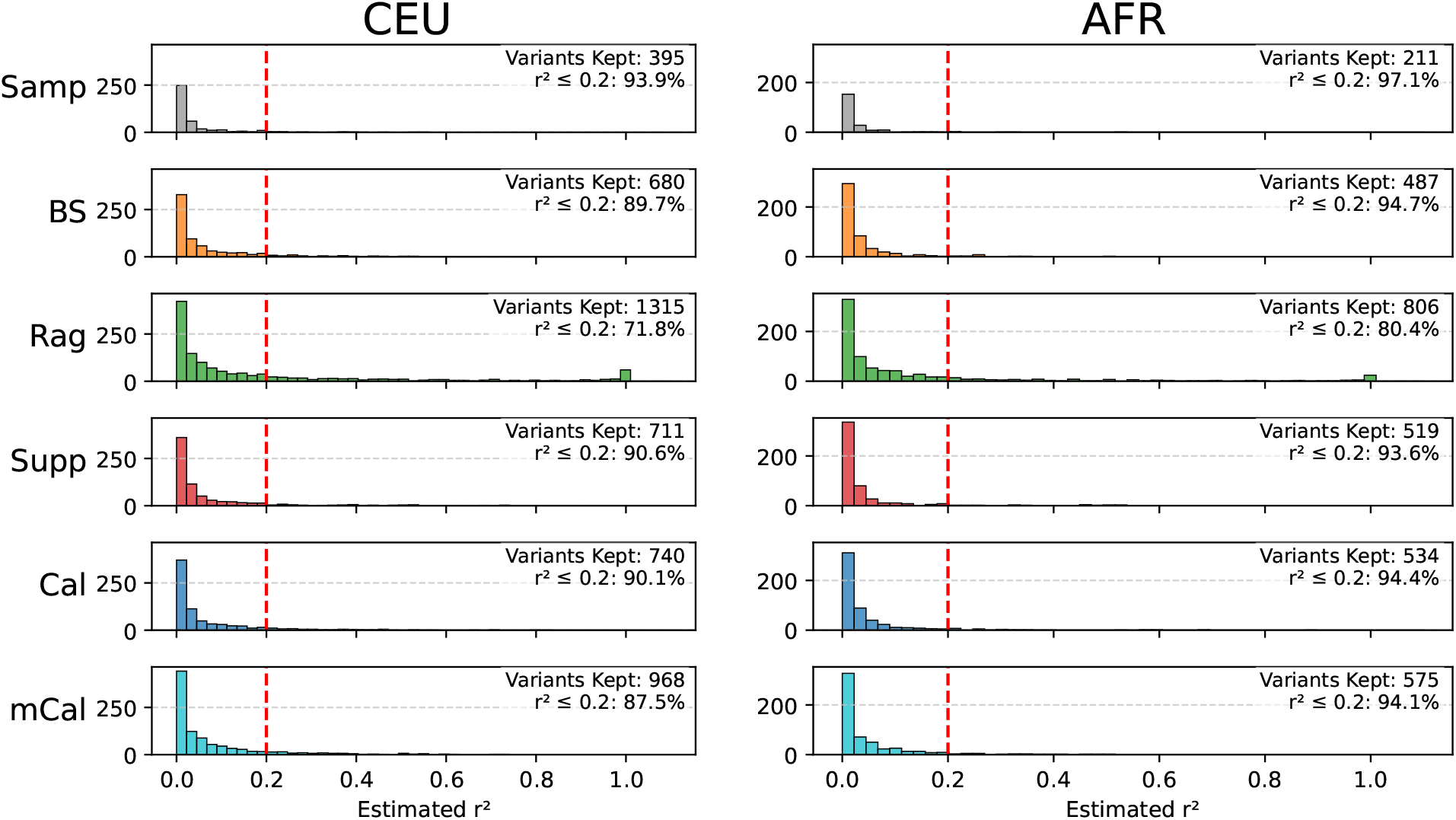
Distribution of *r*^2^ for adjacent SNPs remaining after LD pruning. The analysis was performed on a sample of *n* = 10 individuals. The left panel shows results from the 1000 Genomes CEU population, while the right panel shows results from the simulated AFR population. Each row displays the histogram of the true *r*^2^ values for the set of SNPs kept by each estimator. The red dashed line indicates the pruning threshold (*r*^2^ = 0.2). The annotations in each subplot provide the total number of variants kept and the percentage of remaining adjacent pairs with *r*^2^ ≤ 0.2.

## Discussion

In this work, we present a simulation-based approach to estimate the distribution of the squared correlation and implement a 2-step calibration process to reduce bias in low sample size settings. While simulation studies of population genetics metrics are not novel [12, 26], leveraging modern computational resources allows us to accurately estimate the density of key measures, such as the squared correlation, and use these estimates to calibrate standard estimators. The two-step calibration further corrects for bias, although with a modest increase in variance.

We evaluated the calibration procedure by comparing it with established approaches [5, 11] and a novel estimator introduced in this work. Both real data from the 1000 Genomes Project [14] and simulated data using `stdpopsim` [16] were used to cover diverse scenarios. Across these datasets, the calibrated estimators consistently showed superior performance in terms of RMSE, particularly the 1-step calibration, demonstrating that precise simulation-based density estimates can substantially improve LD metrics in low-sample contexts.

To examine downstream implications, we assessed the classification performance using the F1 score. As with RMSE, the calibrated estimators outperformed other methods, especially for the smallest sample sizes (*n* = 5 and *n* = 10). Analysis of retained variants in pruning experiments further revealed that the calibrated methods strike a better balance between over-pruning and under-pruning, highlighting their practical utility in LD-based analyses.

## Code availability

The implementation of the calibration procedure in `Python` with the code used in the presentation of this work are publicly available in the `GitHub` repository at https://github.com/uliBercovich/SCoLD.

