## Supplementary Material for "Calibration improves estimation of linkage disequilibrium on low sample sizes"

#### S1 First order approximation of Pearson $r^2$

Let's assume that we only observe $G_{is} = H_{is}^m + H_{is}^f$ and $G_{it} = H_{it}^m + H_{it}^f$. Let's assume that $H_{i\cdot}^m$ and $H_{i\cdot}^f$ have a marginal distribution given by a Binomial with parameters $p_\cdot$. While we assume that the pairs $H_{i\cdot}^m$ and $H_{i\cdot}^f$ are independent, we allow some dependence between $H_{is}^m$ with $H_{it}^m$ and $H_{is}^f$ with $H_{it}^f$, denoted by $p_{st}$, the probability of having both reference alleles. Let $D_{st} = p_{st} - p_s p_t$ be the covariance.

Following the method developed by Ragsdale & Gravel [1], where they use unbiased estimators for both the numerator and denominator of Pearson $r^2$, we can study the biases that we introduce by using standard estimators for $D_{st}^2$ and $D_{ss}D_{tt}$. This practice can be theoretically grounded by using the Taylor expansion. Assume strictly positive random variables $X$ and $Y$ with means $\mu_X$ and $\mu_Y$, respectively. Let $f(X,Y) = \frac{X}{Y}$. Then, the first order Taylor expansion at $(\mu_X, \mu_Y)$ is

$$\frac{X}{Y} \approx \frac{\mu_X}{\mu_Y} + (X - \mu_X)\frac{1}{\mu_Y} - (Y - \mu_Y)\frac{\mu_X}{\mu_Y^2}.$$

Hence,

$$\mathbb{E}\left[\frac{X}{Y}\right] \approx \mathbb{E}\left[\frac{\mu_X}{\mu_Y} + (X - \mu_X)\frac{1}{\mu_Y} - (Y - \mu_Y)\frac{\mu_X}{\mu_Y^2}\right] = \frac{\mu_X}{\mu_Y}.$$

Then, it makes sense to study estimators for $r_{st}^2$ given by

$$\widetilde{r_{st}^2} = \frac{\widetilde{D_{st}^2}}{\widetilde{D_{ss}D_{tt}}},$$

where $\widetilde{D_{st}^2}$ and $\widetilde{D_{ss}D_{tt}}$ are unbiased estimators of $D_{st}^2$ and $D_{ss}D_{tt}$, respectively.

Let $\widehat{D}_{st}$ be an estimator of $D_{st}$ given by

$$\widehat{D}_{st} = \frac{1}{2(n-1)}\left(\sum_{i=1}^{n} G_{is}G_{it} - \frac{1}{n}\sum_{i,k} G_{is}G_{kt}\right).$$

The aim is to analytically compute $\mathbb{E}(\widehat{D}_{st}^2)$ and $\mathbb{E}(\widehat{D}_{ss}\widehat{D}_{tt})$ to then calculate the bias terms. To do this, let $\overline{G}_\cdot = \frac{1}{n}\sum_{i=1}^{n} G_{i\cdot}$.

As the computation of this expectations involves summations with up to 4 different indices, a program on the symbolic programming software `SymPy` [2] was developed to facilitate the calculations of the following expectations.

$$\begin{aligned}\mathbb{E}[\widehat{D}_{st}] &= \frac{1}{2(n-1)}\mathbb{E}\left[\left(\sum_i G_{is}G_{it} - \frac{1}{n}\sum_{i,k} G_{is}G_{kt}\right)\right] \\ &= D_{st}.\end{aligned} \tag{S1}$$

$$\begin{aligned}\mathbb{E}[\widehat{D}_{st}\overline{G}_s] &= \frac{1}{2n(n-1)}\mathbb{E}\left[\sum_{i,j} G_{is}G_{js}G_{it} - \frac{1}{n}\sum_{i,j,k} G_{is}G_{js}G_{kt}\right] \\ &= \frac{1}{n}D_{st} + \frac{2(n-1)}{n}D_{st}h_s.\end{aligned} \tag{S2}$$

$$\begin{aligned}\mathbb{E}[\widehat{D}_{st}\overline{G}_s\overline{G}_t] &= \frac{1}{2n^2(n-1)}\mathbb{E}\left[\sum_{i,j,k} G_{is}G_{js}G_{it}G_{kt} - \frac{1}{n}\sum_{i,j,k,\ell} G_{is}G_{js}G_{kt}G_{\ell t}\right] \\ &= \frac{1}{n^2}D_{st} + \frac{2(n-1)}{n^2}D_{st}(h_s+h_t) + \frac{4(n-1)^2}{n^2}D_{st}h_sh_t + \frac{2(n-1)}{n^2}D_{st}^2.\end{aligned} \tag{S3}$$

$$\begin{aligned}\mathbb{E}[\widehat{D}_{ss}\widehat{D}_{tt}] &= \frac{1}{4(n-1)^2}\mathbb{E}\Bigg[\sum_{i,k} G_{is}^2G_{kt}^2 - \frac{1}{n}\sum_{i,k,\ell} G_{is}^2G_{kt}G_{\ell t} - \frac{1}{n}\sum_{i,j,k} G_{is}G_{js}G_{kt}^2 \\ &\qquad + \frac{1}{n^2}\sum_{i,j,k,\ell} G_{is}G_{js}G_{kt}G_{\ell t}\Bigg] \\ &= \frac{1}{2n}D_{st} - \frac{1}{n}D_{st}(h_s+h_t) + \frac{2}{n}D_{st}h_sh_t + \frac{n+1}{n(n-1)}D_{st}^2 + h_s(1-h_s)h_t(1-h_t).\end{aligned} \tag{S4}$$

$$\begin{aligned}\mathbb{E}[\widehat{D}_{st}^2] &= \frac{1}{4(n-1)^2}\mathbb{E}\left[\sum_{i,j} G_{is}G_{js}G_{it}G_{jt} - \frac{2}{n}\sum_{i,j,k} G_{is}G_{js}G_{it}G_{kt} + \frac{1}{n^2}\sum_{i,j,k,\ell} G_{is}G_{js}G_{kt}G_{\ell t}\right] \\ &= \frac{1}{2n}D_{st} - \frac{1}{n}D_{st}(h_s+h_t) + \frac{2}{n}D_{st}h_sh_t + \frac{n^2-n+1}{n(n-1)}D_{st}^2 + \frac{1}{n-1}h_s(1-h_s)h_t(1-h_t).\end{aligned} \tag{S5}$$

Hence, by combining (S1), (S2), (S3), (S4) and (S5), we define

$$\widetilde{D_{st}^2} = \frac{(n-1)^2}{n(n-2)}\widehat{D}_{st}^2 - \frac{n-1}{n(n-2)}\widehat{D}_{ss}\widehat{D}_{tt} - \frac{1}{2(n-1)}\widehat{D}_{st}(\overline{G}_s-1)(\overline{G}_t-1) \tag{S6}$$

and

$$\widetilde{D_{ss}D_{tt}} = -\frac{n-1}{n(n-2)}\widehat{D}_{st}^2 + \frac{(n-1)^2}{n(n-2)}\widehat{D}_{ss}\widehat{D}_{tt} - \frac{1}{2(n-1)}\widehat{D}_{st}(\overline{G}_s-1)(\overline{G}_t-1), \tag{S7}$$

which are unbiased estimators of $D_{st}^2$ and $D_{ss}D_{tt}$ respectively. Then, by using (S6) and (S7), the first order approximation of $\mathbb{E}\left[r_{st}^2\right]$ is given by

$$\widetilde{r_{st}^2} = \frac{\widetilde{D_{st}^2}}{\widetilde{D_{ss}D_{tt}}}. \tag{S8}$$

Then, we developed an estimator based on the Taylor expansion of $r_{st}^2$. In contrast to the similar estimator presented by Ragsdale & Gravel [1], it allows the use of private data as long as we have access to the standard estimates $\widehat{D}$ and $\overline{G}$.

Furthermore, an issue on the aforementioned estimator [1], are the cases where the denominator is 0. The complexity of its formulation makes the study of the roots on the denominator a difficult task. On the other hand, Lemma S1 ensures a formulation of $\widetilde{r^2_{st}}$ where we can grant that the estimator is well defined under some hypothesis.

**Lemma S1.** *Let* $\widetilde{r^2_{st}}$ *be defined as in* (S8). *Then, the estimator can be written as*

$$\widetilde{r^2_{st}} = 1 + \frac{\widehat{D}^2_{st} - \widehat{D}_{ss}\widehat{D}_{tt}}{-\frac{1}{n}\widehat{D}^2_{st} + \frac{n-1}{n}\widehat{D}_{ss}\widehat{D}_{tt} - \frac{n-2}{2(n-1)^2}\widehat{D}_{st}(\overline{G}_s - 1)(\overline{G}_t - 1)}. \tag{S9}$$

*Furthermore, for* $n > 2$ *and* $\widehat{D}_{ss}\widehat{D}_{tt} > \frac{n^2}{4(n-1)^4}$,

$$-\frac{1}{n}\widehat{D}^2_{st} + \frac{n-1}{n}\widehat{D}_{ss}\widehat{D}_{tt} - \frac{n-2}{2(n-1)^2}\widehat{D}_{st}(\overline{G}_s - 1)(\overline{G}_t - 1) > 0,$$

*which ensures that an estimate will be well defined under this assumption.*

*Proof.* By using that $\widetilde{D^2_{st}}$ and $\widetilde{D_{ss}D_{tt}}$ are given by (S6) and (S7), simple algebra shows that $\widetilde{r^2_{st}}$ can be written as in (S9).

For the second part of the lemma, we divide the proof by looking the terms separately. We begin by seeing that

$$\hat{r}^2 = \frac{\widehat{D}^2_{st}}{\widehat{D}_{ss}\widehat{D}_{tt}} \iff \widehat{D}^2_{st} = \hat{r}^2\widehat{D}_{ss}\widehat{D}_{tt}. \tag{S10}$$

Hence,

$$\begin{aligned} \frac{n-1}{n}\widehat{D}_{ss}\widehat{D}_{tt} - \frac{1}{n}\widehat{D}^2_{st} &= \frac{n-1}{n}\widehat{D}_{ss}\widehat{D}_{tt} - \frac{1}{n}\hat{r}^2\widehat{D}_{ss}\widehat{D}_{tt} & \text{(S11)} \\ &> \frac{n-2}{n}\widehat{D}_{ss}\widehat{D}_{tt}. & \text{(S12)} \end{aligned}$$

By using again (S10), we have the known bound that relates the covariance with variances given by

$$\widehat{D}_{st} < \sqrt{\widehat{D}_{ss}\widehat{D}_{tt}}.$$

This implies that

$$\begin{aligned} \frac{n-2}{2(n-1)^2}\widehat{D}_{st}(\overline{G}_s - 1)(\overline{G}_t - 1) &< \frac{n-2}{2(n-1)^2}\sqrt{\widehat{D}_{ss}\widehat{D}_{tt}}(\overline{G}_s - 1)(\overline{G}_t - 1) & \text{(S13)} \\ &< \frac{n-2}{2(n-1)^2}\sqrt{\widehat{D}_{ss}\widehat{D}_{tt}}. & \text{(S14)} \end{aligned}$$

By using the bounds on (S11) and (S13), we get that

$$\begin{aligned} &-\frac{1}{n}\widehat{D}^2_{st} + \frac{n-1}{n}\widehat{D}_{ss}\widehat{D}_{tt} - \frac{n-2}{2(n-1)^2}\widehat{D}_{st}(\overline{G}_s - 1)(\overline{G}_t - 1) \\ &> \frac{n-2}{n}\widehat{D}_{ss}\widehat{D}_{tt} - \frac{n-2}{2(n-1)^2}\sqrt{\widehat{D}_{ss}\widehat{D}_{tt}} \\ &> 0, \end{aligned}$$

where the last bound is given by using that

$$\widehat{D}_{ss}\widehat{D}_{tt} > \frac{n^2}{4(n-1)^4}.$$

□

The hypothesis of $\widehat{D}_{ss}\widehat{D}_{tt} > \frac{n^2}{4(n-1)^4}$ is easily satisfied considering that genotype data is usually preprocessed by using a minor allele frequency filter for extremely rare variants. Hence, both $\widehat{D}_{ss}$ and $\widehat{D}_{tt}$ are not allowed to be too close to zero.

### S2 Calibration of sample-size aware methods

The calibration procedure presented in Equation 2 can be used for already existing methods that do a sample-size correction. To have a more fair comparison, we added a hard bound on each method, so their estimates are always between 0 and 1. Even compared to the bounded methods, their calibrated ones mostly outperforms in terms of accuracy. In particular, this improvement can be seen in the lower sample sizes. In Figure S1 we can see how calibrated methods, noted with (method) "Cal" achieve lower Root Mean Square Error (RMSE) than the hard bounded versions, noted (method) "HB". Moreover, we can see a comparable or even improved performance against the pure "Cal", that was had consistent best accuracy in main Figure 2. This consistent improvement by the calibration the estimators can also be observed in Figure S2. In particular, it is worth noticing that the calibrated version of the Bulik-Sullivan approach [3] appears to have the greatest performance for low $n$. However, we are using for the thorough analysis in the main comparisons the basic sample calibration. This is to avoid p-hacking by selecting the top performer, as we are comparing many alternatives, but it is worth mentioning that there might be an even better method by calibrating sample-size-aware estimates.

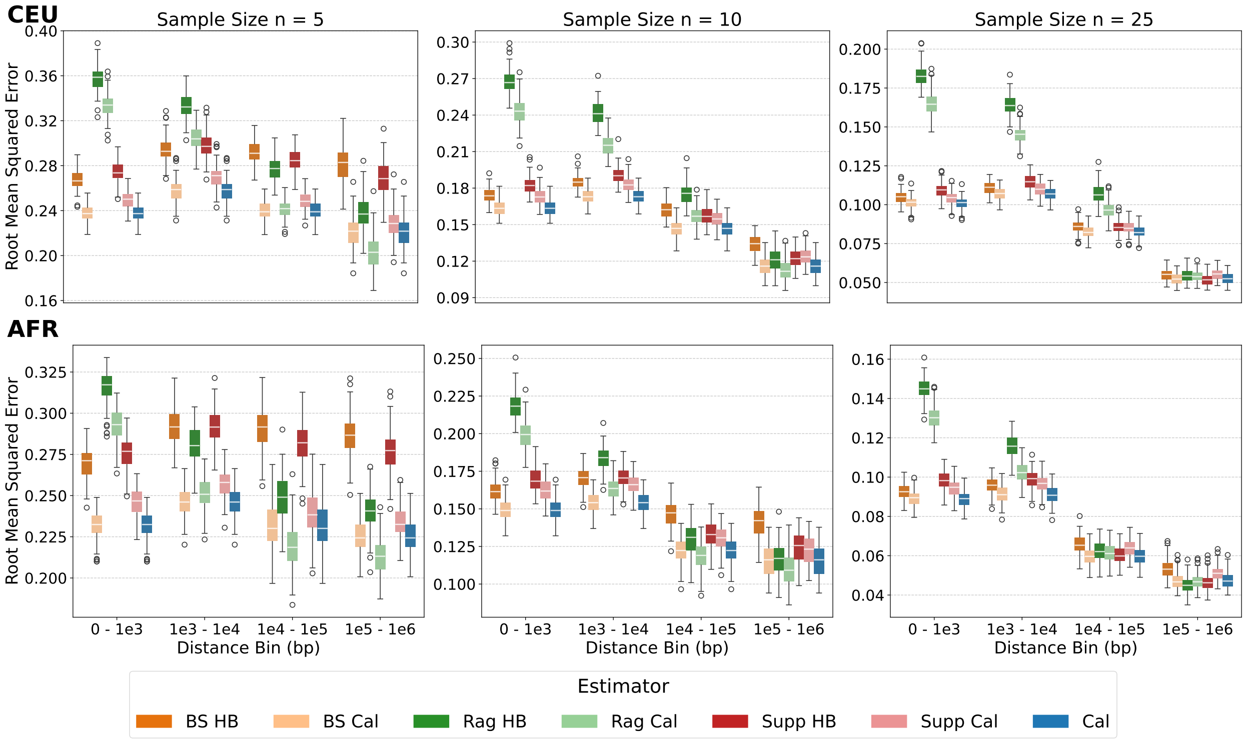


Figure S1: Distribution of Root Mean Squared Error (RMSE) for $r^2$ estimators across genomic distance and sample size. Each row is computed by using different data (CEU and AFR), while each of the three columns corresponds to a different subsample size ($n$). The x-axis in each plot represents genomic distance, divided into bins. Within each bin, the boxplots show the distribution of RMSE values obtained from 100 independent bootstrap replicates, where each color represents a different $r^2$ estimator.

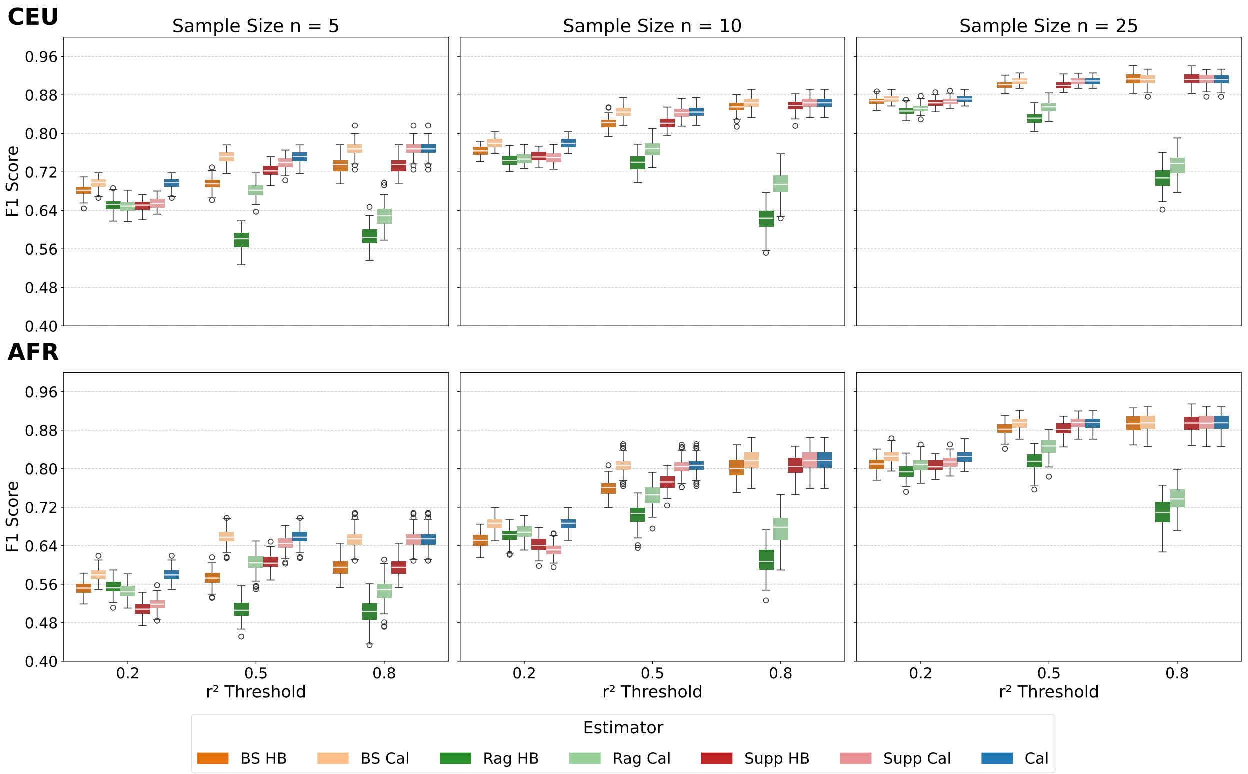


Figure S2: F1 Score for LD pruning performance, compared across populations and sample sizes. The top and bottom rows show performance on data from the CEU and AFR populations, respectively. Each column displays results for a different subsample size ($n$). The x-axis represents the $r^2$ threshold for pruning, while the y-axis shows the F1 score, where a higher value indicates a better balance between precision (avoiding over-pruning) and recall (avoiding under-pruning). The boxplots are based on the resulting F1 score for each method over 100 independent bootstrap replicates.

#### S3 Computation of LD Scores

To further investigate the implications of having more accurate estimates of $r^2$, we measured the Root Mean Square Error (RMSE) of each method when computing LD scores. The LD score of a variant summarizes the amount of genetic variation it tags by aggregating its correlations ($r^2$) with neighboring variants [3]. For a variant $j$, the LD score defined as

$$\ell_j := \sum_{i \in I(j)} r_{ij}^2,$$

where $I(j)$ is the set of variants within a 500,000 bp window centered on variant $j$. To estimate the RMSE, we subsampled our datasets with $n = 5, 10, 25$ individuals. From each subsample, we randomly selected 1000 variants, estimated their LD scores, and compared them to the "true" scores obtained using the full dataset. This procedure was repeated 50 times on independent subsamples and target variants, and the RMSE was computed for each method.

Figure S3 presents the boxplots of the RMSE over the 50 replicates. As expected, the one-step calibration method "Cal" does not yield accurate LD score estimates. This is due to the fact that, by only admitting values between 0 and 1, the upward bias, although heavily reduced, is still there. This issue is particularly relevant in metrics where the total LD score is more important than the accuracy of individual pairwise estimates, i.e., when bias dominates over variance. In contrast, the sample-aware estimators ("BS", "Rag", and "Supp") perform substantially better, with "BS" often showing a slight advantage over the others. The best performance, however, is consistently achieved by the two-step calibration method ("mCal"), which outperforms all other approaches, especially at smaller sample sizes ($n = 5$ and $n = 10$). This improvement arises because "mCal" trades a small loss in accuracy on dependent variant pairs for a much stronger reduction in bias on independent pairs, leading to overall more reliable LD score estimates.

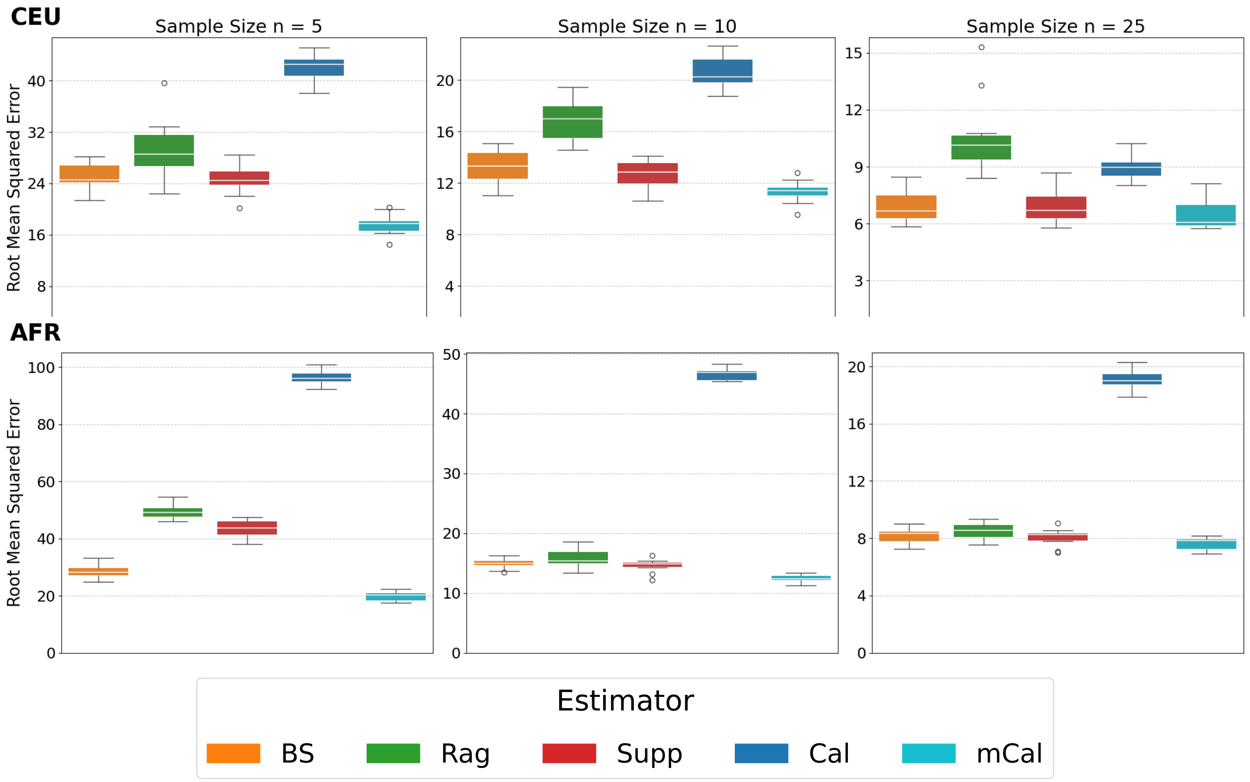


Figure S3: Root Mean Squared Error (RMSE) for LD score estimation, compared across populations and sample sizes. The top and bottom rows show results for the CEU and AFR populations, respectively. Each panel corresponds to a different subsample size ($n$). The y-axis shows the RMSE of the estimated LD scores relative to the true values obtained from the full sample. Each RMSE value is computed over the LD scores of 1,000 randomly chosen SNPs. The boxplots summarize results across 50 independent replicates, each based on random subsamples of individuals and variants.

### Supplementary Algorithms

**Algorithm S1** Estimation of $g_{p_s,p_t}(\rho^2)$

**Input:** Sample size $n$, discretized grid of $\rho^2$ values between 0 and 1 ($\texttt{grid\_}\rho^2$), number of repetitions $\texttt{Nrep}$.
**Output:** Estimated map $\widehat{g}_{(p_s,p_t)}(\rho^2)$.

**for** allele frequencies $(p_s, p_t) \in \{1/(2n), \ldots, (2n-1)/(2n)\}^2$ **do**
  **for** each $\rho^2$ in $\texttt{grid\_}\rho^2$ **do**
    **if** $(p_s, p_t, \rho^2)$ is a valid tuple **then** ▷ satisfies $D$ bounds
      **for** rep $= 1, \ldots, \texttt{Nrep}$ **do**
        **repeat**
          Simulate genotype matrix $G \sim (n, p_s, p_t, \rho^2)$
        **until** $\text{Var}(G_{\cdot s}) > 0$ and $\text{Var}(G_{\cdot t}) > 0$ ▷ $r^2$ has to be well defined
        Compute $r^2_{\texttt{rep}} \leftarrow r^2(G)$ ▷ following Equation (1)
      **end for**
      $\widehat{g}_{(p_s,p_t)}(\rho^2) \leftarrow \frac{1}{\texttt{Nrep}} \sum_{\texttt{rep}=1}^{\texttt{Nrep}} r^2_{\texttt{rep}}$
    **end if**
  **end for**
**end for**

**Algorithm S2** Calibration of observed $r^2$ using $\widehat{g}$

**Input:** Two-locus genotype matrix $G$, precomputed map $\widehat{g}$.
**Output:** Calibrated estimate $\widehat{r^2_{st}}$.

$(\widehat{p}_s, \widehat{p}_t) \leftarrow \frac{1}{2n} \left(\sum_{i=1}^n G_{is}, \sum_{i=1}^n G_{it}\right)$ ▷ estimate allele frequencies
Compute $r^2_{st} \leftarrow r^2(G_{\cdot s}, G_{\cdot t})$ ▷ following Equation (1)
$\widehat{r^2_{st}} \leftarrow \widehat{g}^{-1}_{(\widehat{p}_s,\widehat{p}_t)}(r^2_{st})$ ▷ calibrate via inverse lookup/interpolation

### Supplementary Figures

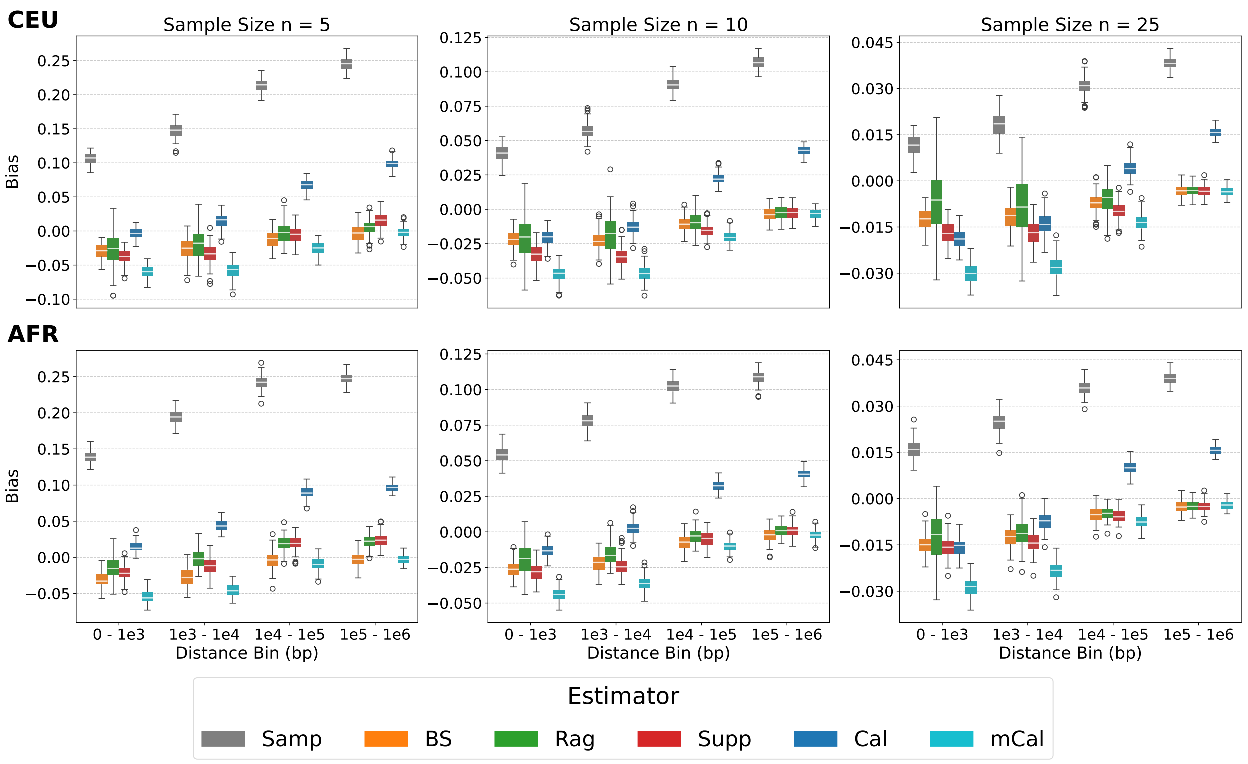


Figure S4: Bias of the estimates as a function of distance and sample size. Each row displays the distributions for different sample sizes. The x-axis shows genomic distance bins. The boxplots within each bin represent the distribution of the bias calculated over 100 bootstrap replicates.

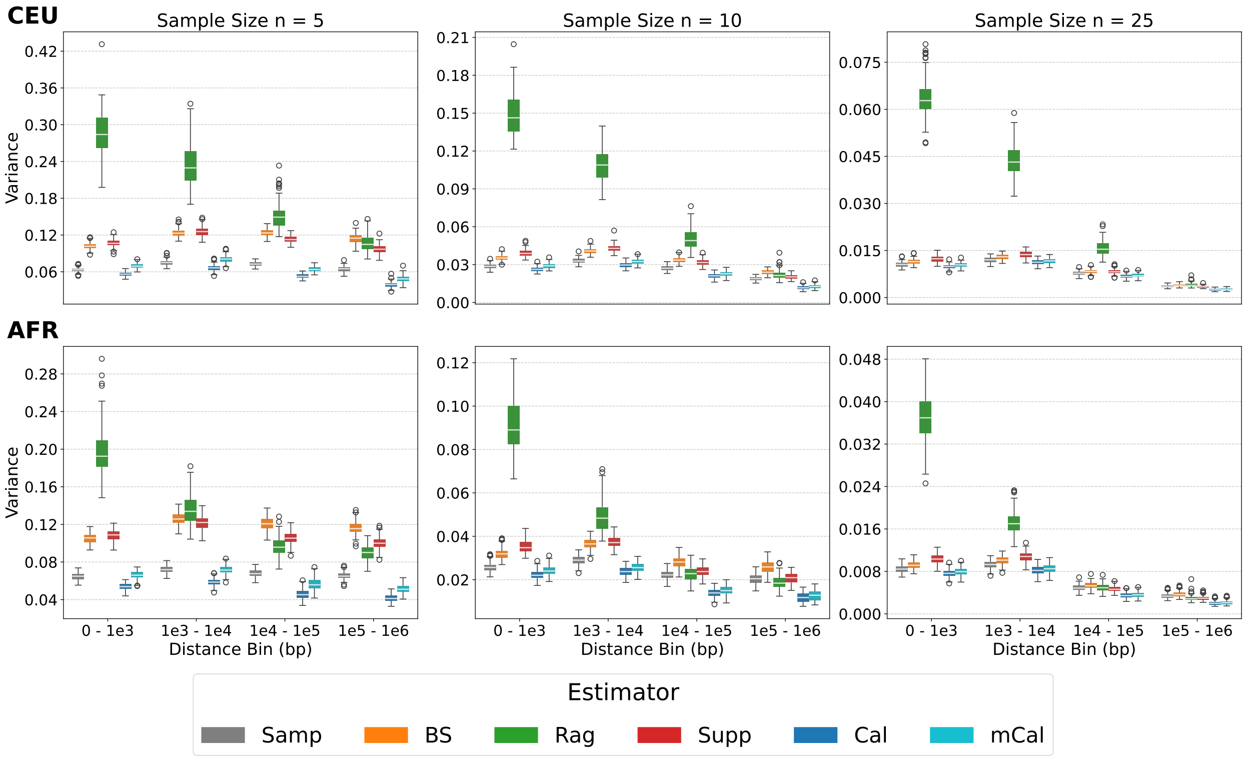


Figure S5: Variances of the errors of the estimates as a function of distance and sample size. Each row displays the distributions for different sample sizes. The x-axis shows genomic distance bins. The boxplots within each bin represent the distribution of the variance of the errors calculated over 100 bootstrap replicates.

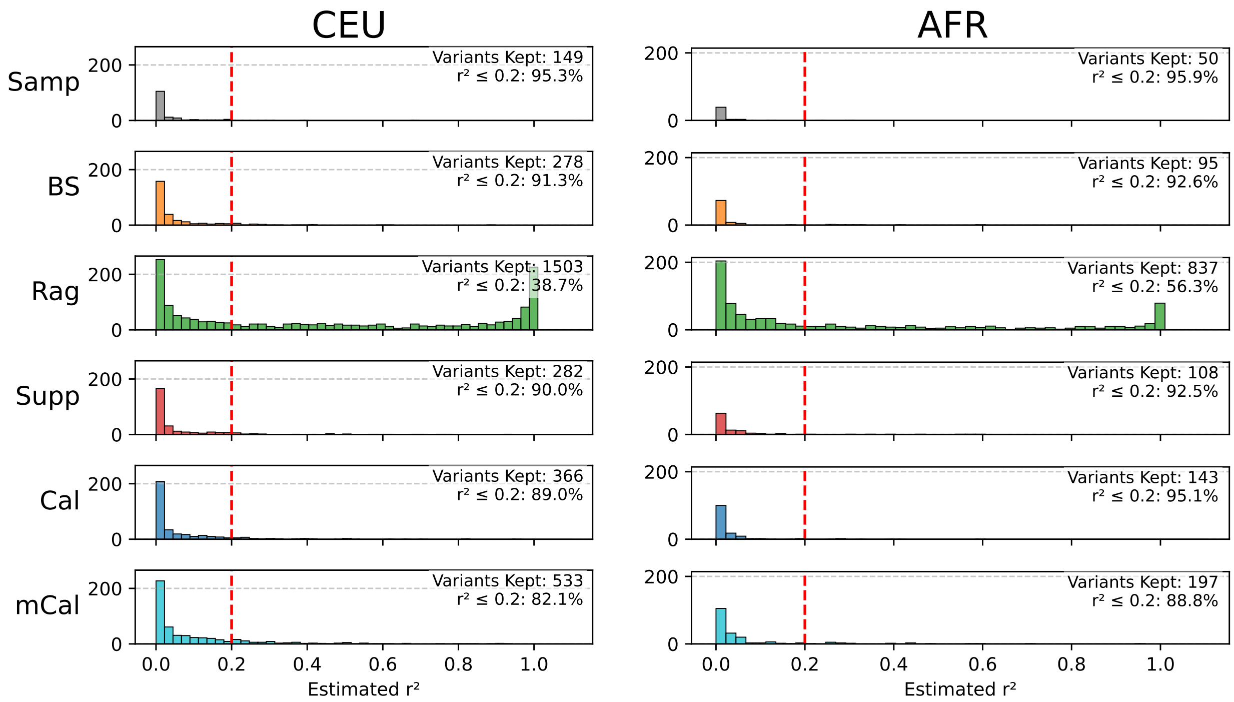


Figure S6: Distribution of $r^2$ for adjacent SNPs remaining after LD pruning. The analysis was performed on a sample of $n = 5$ individuals. The left panel shows results from the 1000 Genomes CEU population, while the right panel shows results from the simulated AFR population. Each row displays the histogram of the true $r^2$ values for the set of SNPs kept by each estimator. The red dashed line indicates the pruning threshold ($r^2 = 0.2$). The annotations in each subplot provide the total number of variants kept and the percentage of remaining adjacent pairs with $r^2 \leq 0.2$.

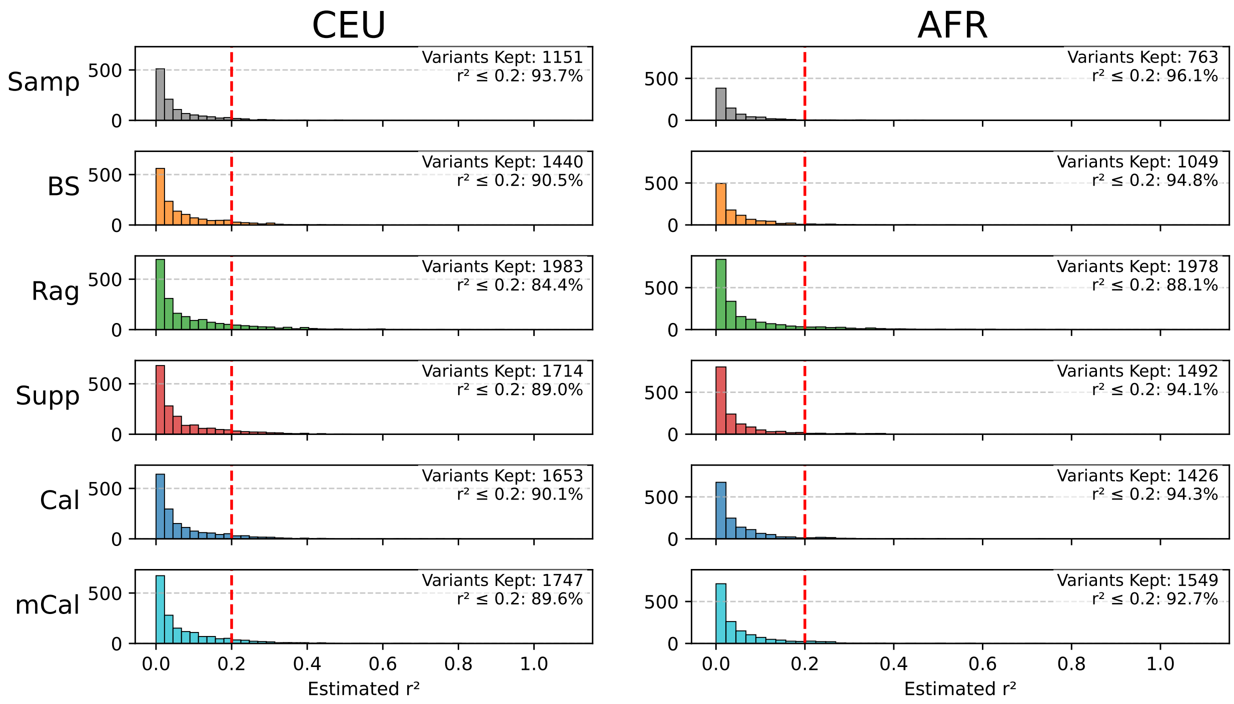


Figure S7: Distribution of $r^2$ for adjacent SNPs remaining after LD pruning. The analysis was performed on a sample of $n = 25$ individuals. The left panel shows results from the 1000 Genomes CEU population, while the right panel shows results from the simulated AFR population. Each row displays the histogram of the true $r^2$ values for the set of SNPs kept by each estimator. The red dashed line indicates the pruning threshold ($r^2 = 0.2$). The annotations in each subplot provide the total number of variants kept and the percentage of remaining adjacent pairs with $r^2 \leq 0.2$.

#### Supplementary Tables

| Bin Range (bp) | Available Pairs |
|---|---:|
| [0, 1,000) | 9,769 |
| [1,000, 10,000) | 73,895 |
| [10,000, 100,000) | 656,239 |
| [[100,000, 1,000,000) | 5,858,331 |

Table S1: Number of available SNP pairs per distance bin. Analysis was performed on the CEU dataset, which corresponds to a genomic region of Chromosome 22 containing 15,938 SNPs from the 1000 Genomes Project.

| Bin Range (bp) | Available Pairs |
|---|---:|
| [0, 1,000) | 20,229 |
| [1,000, 10,000) | 171,386 |
| [10,000, 100,000) | 1,645,073 |
| [[100,000, 1,000,000) | 15,637,545 |

Table S2: Number of available SNP pairs per distance bin. Analysis was performed on the AFR dataset, which corresponds to a genomic region of Chromosome 22 containing 13,543 SNPs from a simulated African population.

| | **Bias** | | **Variance** | | **RMSE** | |
|---|---|---|---|---|---|---|
| **Estimator** | **Mean** | **Std Dev** | **Mean** | **Std Dev** | **Mean** | **Std Dev** |
| **Sample Size n = 5** | | | | | | |
| Samp | $1.78 \times 10^{-1}$ | $5.55 \times 10^{-2}$ | $6.90 \times 10^{-2}$ | $6.56 \times 10^{-3}$ | $3.21 \times 10^{-1}$ | $3.30 \times 10^{-2}$ |
| BS | $-1.77 \times 10^{-2}$ | $1.64 \times 10^{-2}$ | $1.16 \times 10^{-1}$ | $1.11 \times 10^{-2}$ | $3.41 \times 10^{-1}$ | $1.61 \times 10^{-2}$ |
| Rag | $-1.06 \times 10^{-2}$ | $2.32 \times 10^{-2}$ | $1.94 \times 10^{-1}$ | $7.46 \times 10^{-2}$ | $4.33 \times 10^{-1}$ | $8.56 \times 10^{-2}$ |
| Supp | $-1.53 \times 10^{-2}$ | $2.50 \times 10^{-2}$ | $1.11 \times 10^{-1}$ | $1.24 \times 10^{-2}$ | $3.33 \times 10^{-1}$ | $1.89 \times 10^{-2}$ |
| Cal | $4.47 \times 10^{-2}$ | $4.12 \times 10^{-2}$ | $5.38 \times 10^{-2}$ | $1.05 \times 10^{-2}$ | $2.39 \times 10^{-1}$ | $1.65 \times 10^{-2}$ |
| mCal | $-3.50 \times 10^{-2}$ | $2.56 \times 10^{-2}$ | $6.57 \times 10^{-2}$ | $1.26 \times 10^{-2}$ | $2.59 \times 10^{-1}$ | $2.72 \times 10^{-2}$ |
| **Sample Size n = 10** | | | | | | |
| Samp | $7.38 \times 10^{-2}$ | $2.69 \times 10^{-2}$ | $2.69 \times 10^{-2}$ | $5.45 \times 10^{-3}$ | $1.82 \times 10^{-1}$ | $9.89 \times 10^{-3}$ |
| BS | $-1.49 \times 10^{-2}$ | $1.01 \times 10^{-2}$ | $3.34 \times 10^{-2}$ | $6.49 \times 10^{-3}$ | $1.83 \times 10^{-1}$ | $1.87 \times 10^{-2}$ |
| Rag | $-1.24 \times 10^{-2}$ | $1.39 \times 10^{-2}$ | $8.25 \times 10^{-2}$ | $5.08 \times 10^{-2}$ | $2.72 \times 10^{-1}$ | $9.42 \times 10^{-2}$ |
| Supp | $-2.15 \times 10^{-2}$ | $1.46 \times 10^{-2}$ | $3.36 \times 10^{-2}$ | $9.09 \times 10^{-3}$ | $1.83 \times 10^{-1}$ | $2.71 \times 10^{-2}$ |
| Cal | $7.78 \times 10^{-3}$ | $2.63 \times 10^{-2}$ | $2.23 \times 10^{-2}$ | $7.11 \times 10^{-3}$ | $1.50 \times 10^{-1}$ | $2.27 \times 10^{-2}$ |
| mCal | $-2.96 \times 10^{-2}$ | $1.92 \times 10^{-2}$ | $2.43 \times 10^{-2}$ | $7.81 \times 10^{-3}$ | $1.57 \times 10^{-1}$ | $2.92 \times 10^{-2}$ |
| **Sample Size n = 25** | | | | | | |
| Samp | $2.48 \times 10^{-2}$ | $1.09 \times 10^{-2}$ | $8.44 \times 10^{-3}$ | $3.28 \times 10^{-3}$ | $9.45 \times 10^{-2}$ | $1.55 \times 10^{-2}$ |
| BS | $-8.54 \times 10^{-3}$ | $4.97 \times 10^{-3}$ | $9.13 \times 10^{-3}$ | $3.54 \times 10^{-3}$ | $9.39 \times 10^{-2}$ | $2.04 \times 10^{-2}$ |
| Rag | $-5.89 \times 10^{-3}$ | $7.77 \times 10^{-3}$ | $3.18 \times 10^{-2}$ | $2.39 \times 10^{-2}$ | $1.63 \times 10^{-1}$ | $7.40 \times 10^{-2}$ |
| Supp | $-1.18 \times 10^{-2}$ | $6.58 \times 10^{-3}$ | $9.44 \times 10^{-3}$ | $4.06 \times 10^{-3}$ | $9.52 \times 10^{-2}$ | $2.34 \times 10^{-2}$ |
| Cal | $-3.28 \times 10^{-3}$ | $1.43 \times 10^{-2}$ | $7.60 \times 10^{-3}$ | $3.41 \times 10^{-3}$ | $8.57 \times 10^{-2}$ | $2.15 \times 10^{-2}$ |
| mCal | $-1.88 \times 10^{-2}$ | $1.14 \times 10^{-2}$ | $7.90 \times 10^{-3}$ | $3.56 \times 10^{-3}$ | $8.84 \times 10^{-2}$ | $2.39 \times 10^{-2}$ |

Table S3: Performance metrics (mean, standard deviation and RMSE) for $r^2$ estimators across different sample sizes in the CEU dataset.

| | Bias | | Variance | | RMSE | |
|---|---|---|---|---|---|---|
| **Estimator** | **Mean** | **Std Dev** | **Mean** | **Std Dev** | **Mean** | **Std Dev** |
| **Sample Size n = 5** | | | | | | |
| Samp | $2.06 \times 10^{-1}$ | $4.43 \times 10^{-2}$ | $6.75 \times 10^{-2}$ | $4.92 \times 10^{-3}$ | $3.33 \times 10^{-1}$ | $2.87 \times 10^{-2}$ |
| BS | $-1.65 \times 10^{-2}$ | $1.72 \times 10^{-2}$ | $1.17 \times 10^{-1}$ | $1.02 \times 10^{-2}$ | $3.43 \times 10^{-1}$ | $1.45 \times 10^{-2}$ |
| Rag | $6.58 \times 10^{-3}$ | $1.92 \times 10^{-2}$ | $1.30 \times 10^{-1}$ | $4.57 \times 10^{-2}$ | $3.56 \times 10^{-1}$ | $6.04 \times 10^{-2}$ |
| Supp | $2.54 \times 10^{-3}$ | $2.22 \times 10^{-2}$ | $1.09 \times 10^{-1}$ | $1.02 \times 10^{-2}$ | $3.30 \times 10^{-1}$ | $1.53 \times 10^{-2}$ |
| Cal | $6.12 \times 10^{-2}$ | $3.43 \times 10^{-2}$ | $5.00 \times 10^{-2}$ | $7.88 \times 10^{-3}$ | $2.34 \times 10^{-1}$ | $1.33 \times 10^{-2}$ |
| mCal | $-2.68 \times 10^{-2}$ | $2.39 \times 10^{-2}$ | $6.13 \times 10^{-2}$ | $9.42 \times 10^{-3}$ | $2.49 \times 10^{-1}$ | $2.06 \times 10^{-2}$ |
| **Sample Size n = 10** | | | | | | |
| Samp | $8.59 \times 10^{-2}$ | $2.21 \times 10^{-2}$ | $2.43 \times 10^{-2}$ | $3.99 \times 10^{-3}$ | $1.79 \times 10^{-1}$ | $1.00 \times 10^{-2}$ |
| BS | $-1.43 \times 10^{-2}$ | $1.13 \times 10^{-2}$ | $3.06 \times 10^{-2}$ | $4.89 \times 10^{-3}$ | $1.75 \times 10^{-1}$ | $1.44 \times 10^{-2}$ |
| Rag | $-8.89 \times 10^{-3}$ | $1.14 \times 10^{-2}$ | $4.56 \times 10^{-2}$ | $3.00 \times 10^{-2}$ | $2.03 \times 10^{-1}$ | $6.78 \times 10^{-2}$ |
| Supp | $-1.40 \times 10^{-2}$ | $1.34 \times 10^{-2}$ | $2.93 \times 10^{-2}$ | $7.67 \times 10^{-3}$ | $1.71 \times 10^{-1}$ | $2.35 \times 10^{-2}$ |
| Cal | $1.56 \times 10^{-2}$ | $2.23 \times 10^{-2}$ | $1.79 \times 10^{-2}$ | $5.60 \times 10^{-3}$ | $1.35 \times 10^{-1}$ | $1.89 \times 10^{-2}$ |
| mCal | $-2.36 \times 10^{-2}$ | $1.80 \times 10^{-2}$ | $1.94 \times 10^{-2}$ | $6.05 \times 10^{-3}$ | $1.40 \times 10^{-1}$ | $2.44 \times 10^{-2}$ |
| **Sample Size n = 25** | | | | | | |
| Samp | $2.90 \times 10^{-2}$ | $9.62 \times 10^{-3}$ | $6.50 \times 10^{-3}$ | $2.52 \times 10^{-3}$ | $8.53 \times 10^{-2}$ | $1.24 \times 10^{-2}$ |
| BS | $-8.73 \times 10^{-3}$ | $5.85 \times 10^{-3}$ | $7.06 \times 10^{-3}$ | $2.73 \times 10^{-3}$ | $8.29 \times 10^{-2}$ | $1.72 \times 10^{-2}$ |
| Rag | $-7.53 \times 10^{-3}$ | $6.16 \times 10^{-3}$ | $1.55 \times 10^{-2}$ | $1.38 \times 10^{-2}$ | $1.12 \times 10^{-1}$ | $5.52 \times 10^{-2}$ |
| Supp | $-9.53 \times 10^{-3}$ | $6.35 \times 10^{-3}$ | $7.20 \times 10^{-3}$ | $3.50 \times 10^{-3}$ | $8.27 \times 10^{-2}$ | $2.21 \times 10^{-2}$ |
| Cal | $7.49 \times 10^{-4}$ | $1.29 \times 10^{-2}$ | $5.34 \times 10^{-3}$ | $2.71 \times 10^{-3}$ | $7.17 \times 10^{-2}$ | $1.91 \times 10^{-2}$ |
| mCal | $-1.54 \times 10^{-2}$ | $1.13 \times 10^{-2}$ | $5.54 \times 10^{-3}$ | $2.80 \times 10^{-3}$ | $7.38 \times 10^{-2}$ | $2.16 \times 10^{-2}$ |

Table S4: Performance metrics (mean, standard deviation and RMSE) for $r^2$ estimators across different sample sizes in the AFR dataset.
